# Single cell multi-omic whole genome sequencing and chromatin accessibility profiling reveals genome-epigenome coevolution in colorectal cancer

**DOI:** 10.1101/2025.04.15.648959

**Authors:** Maximilian Mossner, Chloé Colson, Qingli Guo, Freddie Whiting, Heather Grant, Amanda Stafford, Peter Kyle, Daniella Donato-Brown, Jamie Murphy, Andrea Sottoriva, Ann-Marie Baker, Trevor A Graham

## Abstract

Epigenetic alterations co-evolve with genetic mutations to drive carcinogenesis and treatment response. Resolving genome-epigenome coevolution requires accurate multi-omic single cell measurement. Here we develop a new technology called “double ATAC” (dATAC) for high-throughput, high-quality, concurrent whole genome sequencing and chromatin accessibility profiling of individual somatic cells. dATAC is a “one pot” method that uses two rounds of tagmentation to sequentially label open chromatin regions and then the whole genome, and produces data of the same quality as current leading single-omic methods. Using colorectal cancer as a model system, we apply dATAC to reveal convergent reorganisation of the epigenome across expanding drug-resistant clones during 5-FU chemotherapy exposure, remarkable stoichiometry of chromatin accessibility at somatic copy number alterations, and the clonal expansion of copy-number altered T cells in the stroma of metastatic disease. dATAC is a robust single cell technology to accurately profile genome-epigenome coevolution across tissues, diseases and species.

## Introduction

Cancers evolve through natural selection acting upon heterogeneity that exists within the cancer cell population. The role of genetic heterogeneity in driving cancer evolution is well-established^1–5^, including at single cell resolution^6–10^, but the contribution of epigenetic heterogeneity is less well understood. Changes to the epigenome could themselves be drivers of somatic evolution, or they could be passenger events that clonally expand due to a driving genetic alteration in a clone. There are also mechanistic links between genetic mutation and epigenetic alteration (“epimutation”). For example, mutations accrue at a higher rate in regions of closed chromatin^11^, and somatic mutations may precipitate epigenetic alterations^12, 13^. The ability to delineate the role of the epigenome in driving cancer evolution, and its interrelationship with genetic evolution, has been limited by the availability of methods to simultaneously, and accurately, profile the genome and epigenome at single cell resolution.

Here, we have developed “dATAC” (<u>d</u>ouble <u>A</u>ssay for <u>T</u>ransposase-<u>A</u>ccessible <u>C</u>hromatin), a high-throughput high-resolution method for co-measurement of whole genome and chromatin accessibility profiles in single nuclei. dATAC utilises two rounds of tagmentation: the first round is performed in intact nuclei to tagment accessible chromatin with an “A” barcode adapter, followed by single cell isolation, complete nucleosome disruption and whole genome tagmentation with a second “B” barcode adapter. Single cells are then barcoded, pooled and sequenced, and then A and B tagment barcodes are used to reconstruct chromatin architecture. dATAC builds upon prior work: Queitsch and colleagues previously reported a combined whole genome sequencing (WGS) and chromatin accessibility (ATAC-seq) method utilising the popular 10X Genomics single cell platform^14^, however data quality is compromised because of the requirement for fixed and intact nuclei throughout the protocol. Single nucleus ATACseq (snATAC-seq) has been combined with targeted capture and sequencing of single nucleotide variants (SNVs) hotspots and applied to study tumour evolution in haematologic neoplasms^15, 16^.

Here we applied dATAC methodology to study genome-epigenome coevolution in colorectal cancer (CRC). In a CRC cell line treated with chemotherapy, dATAC revealed convergent and parallel epigenome evolution in distinct tumour clones, allowing us to map the interrelationship between copy number alteration and changes to chromatin accessibility. In patient samples, dATAC identified a clonally expanded T cell population bearing a somatic copy number alteration in the stroma of metastatic disease.

## Results

### dATAC technology for combined WGS and ATAC profiling in single cells

We constructed “double ATAC” (dATAC), a single nucleus sequencing protocol for very high quality co-measurement of chromatin accessibility and whole genome sequence (Fig. 1). dATAC utilises S3-chemistry by Mulqueen *et al* which introduces indexing on the level of tagmentation and improves the conversion rate of tagmented DNA into functional NGS library fragments^17^. The dATAC protocol is further simplified and optimised to enable robust “one-pot” single nucleus multi-omics in 384-well plate format. First, open chromatin tagmentation is performed in bulk intact nuclei using a Tn5 enzyme loaded with a custom barcoded adapter (barcode A). Single nuclei are then sorted individually into 384-well plates containing lysis buffer. After complete lysis of nuclear and histone proteins in the plate, the previously closed chromatin is fully exposed (accessible) and subjected to a second round of tagmentation using Tn5 enzyme loaded with a second distinct custom barcoded adapter (barcode B). Individual single cell libraries are then PCR amplified using indexing primers, pooled and sequenced. Subsequently, fragments derived from open or closed chromatin can be bioinformatically separated using the inline barcodes A and B as a readout for accessible chromatin and whole genome respectively.

**Figure 1:**
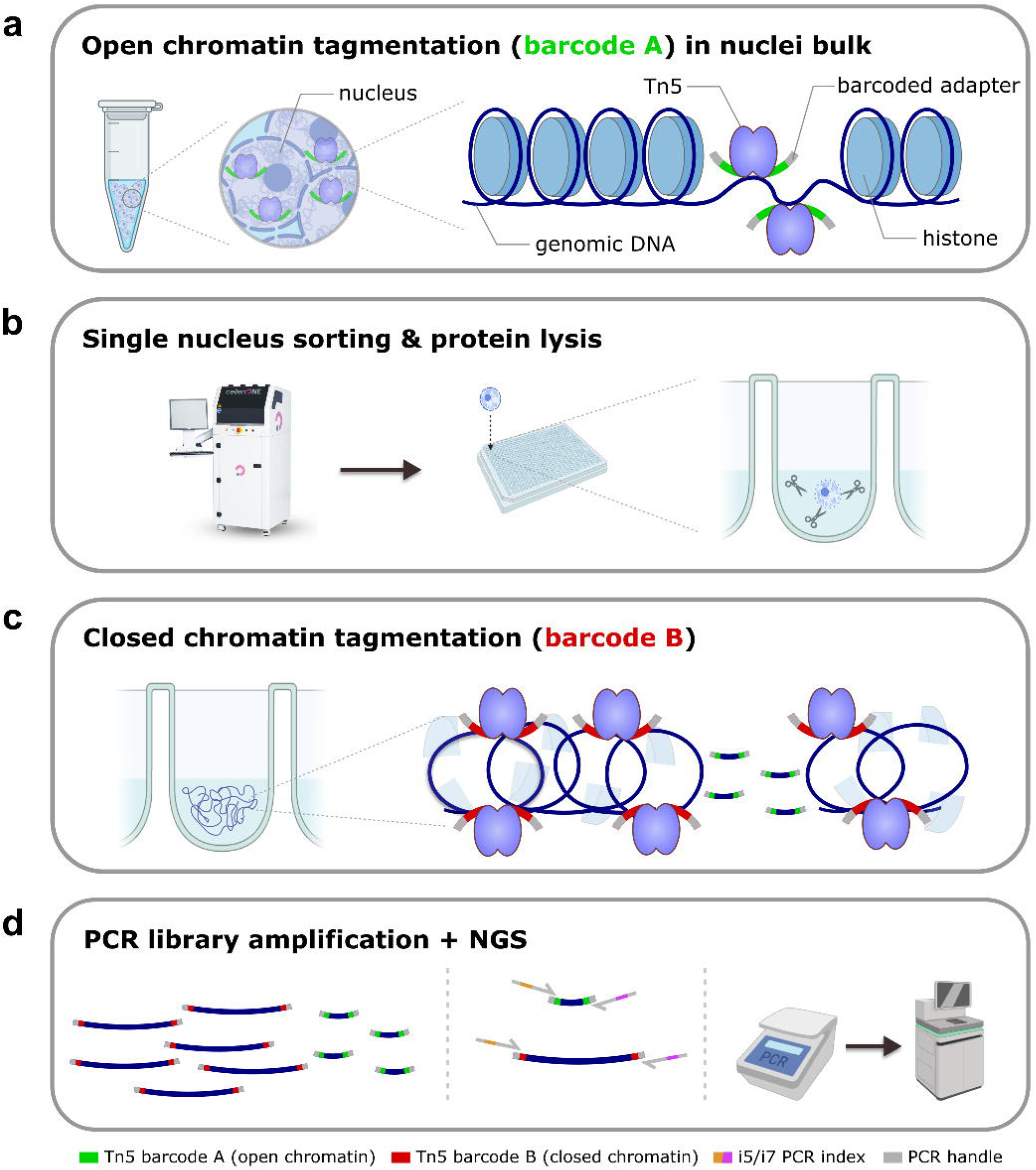
Schematic of dATAC workflow. (a) Tagmentation is initially performed in a nuclei suspension with intact chromatin structure, so that only open chromatin regions are tagmented. The first round tagment enzyme is loaded with an “A” barcode. (b) Nuclei are dispensed into a plate with individual barcode-indexed wells and protein is digested. (c) A second round of tagmentation is performed with an enzyme loaded with a “B” barcode. As chromatin has been digested, the second-round tagmentation is uniform across the genome. (d) Libraries are pooled and sequenced, and cell identity and “A” and “B” barcodes are recovered in the downstream bioinformatics pipeline. Figure was prepared with images from Biorender.

### dATAC enables sensitive co-measurement of chromatin accessibility and whole-genome DNA profiles in single cells

We first evaluated the performance of the open chromatin measurement by dATAC in cancer cell lines. We performed only the first part of the dATAC protocol, labelling open chromatin with barcode A, and skipping the second whole genome tagmentation step (no barcode B). Libraries had high complexity, with median unique fragment counts per nucleus of 121,610, 97,075 for H1299 and MCF7 cells and on average 127,490 for SW620 cells that were either untreated or 5-fluorouracil (5-FU) treated (range 96,317-225,105, T0a/b: untreated control, T1a/b: 5-FU short burst treatment over 5 days) (Extended Data Fig. 1a-d), comparable to current gold standard methods for snATAC analysis^17–19^. Transcription start site (TSS) enrichment was high for all cell lines (mean 9.00, range 6.414-11.703, Extended Data Fig. 1a) and fragment size profiles showed the characteristic nucleosome banding distribution for ATAC preparations (Extended Data Fig. 1b). Uniform manifold approximation and projection (UMAP) clustering of chromatin accessibility data for all tested cell lines showed clear separation based on cell type and treatment condition (Extended Data Fig. 1e). This was further corroborated by the presence of highly cell type specific peak profiles as demonstrated for select genomic loci (Extended Data Fig 1f-g). Overall, these data indicate that chromatin accessibility profiling within dATAC produces high quality data that accurately recovers cell line-specific epigenetic profiles.

We next evaluated multi-modal dATAC performance by comparing WGS data from dATAC processed cells to WGS data generated by “Direct Library Prep plus” (DLP+) a widely-used single-omics WGS assay^20^. DLP+ was adapted for use in a 384-well plate format (see methods). DLP+ single-omics WGS was performed on n=384 cells and dATAC multi-omics analysis on 192 nuclei from SW620 untreated control cells (“SW620 T0”), of which n=280 (DLP) and n=182 (dATAC) generated a minimum of 1 million usable reads (combining A and B barcodes for dATAC), with an average of 8.5 x 10^6 reads per cell (standard deviation (S.D.): 4.7 x 10^6) for DLP+ and 9.9 x 10^6 (S.D.: 4.7 x 10^6) reads per nucleus for dATAC. DLP and dATAC assay copy number alteration (CNA) measurements showed concordant WGS karyotype profiles that were characteristic for the SW620 cell line (Fig. 2a-b) and generally produced high-resolution single cell CNA profiles (Fig. 2c-d). However, the complexity of dATAC libraries was significantly higher than for DLP+ libraries (dATAC average duplication rate of 28% (range 22-30%) for five independent dATAC libraries compared to an average duplication rate of 53% (range: 51-56%) in three DLP+ libraries at a sequencing depth of ∼10 million mapped reads/cell) (Fig. 2e).

**Figure 2.**
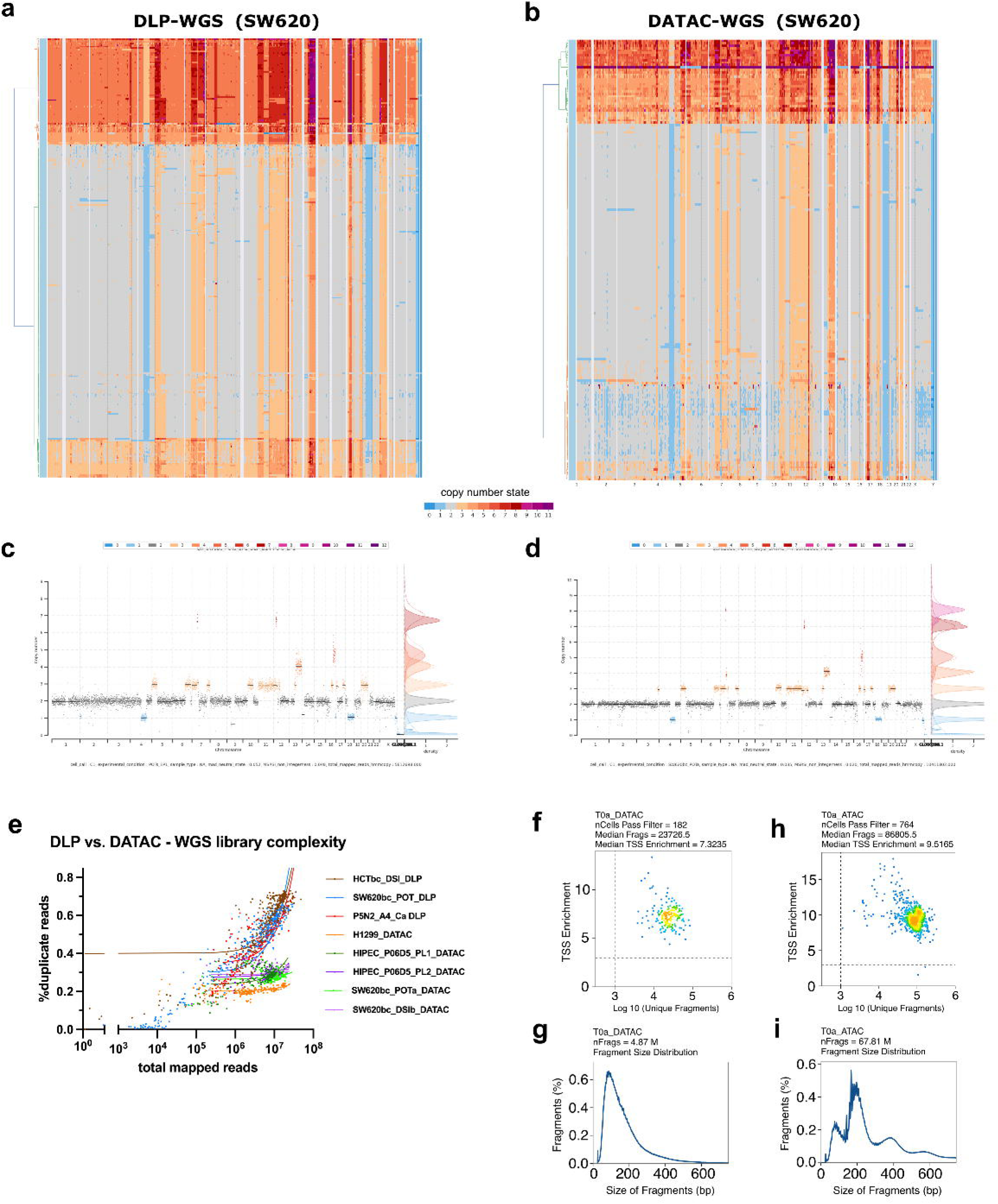
Technical performance of dATAC assay. Single cell whole genome sequencing from SW620 colorectal cancer cells were generated using (a) the single-omics Direct Library Preparation (DLP+) and (b) multi-omic dATAC assay with highly comparable results. (c-d) Individual exemplary single cell karyotypes profiles indicated high resolution for detection of chromosome-level and micro-lesions using both the DLP+ (c) and dATAC assay (d). (e-f) The dATAC assay was run in “single cell ATAC-only” mode (i.e. skipping the second tagmentation) and produced high complexity single cell libraries with high transcription start site (TSS) enrichment, high median number of reads per cell (e) and characteristic fragment size profiles (f) on par with current gold standard single cell ATAC approaches. (g) dATAC assay preparation in “multi-omics” mode (two consecutive tagmentations) confirmed high TSS enrichment and read numbers per cell when analysing barcode “A”/open chromatin-derived reads. (h) The fragment size distribution of the A barcode (ATAC round) in dATAC processed cells shows a shorter fragment size likely due to re-tagmentation of open-chromatin fragments with the B barcode. (i) Whole genome sequencing (WGS) library complexity was considerably higher in for dATAC processed cells compared to single-omic DLP+ processing.

In dATAC processed cells, co-measurement of chromatin accessibility revealed high numbers of unique fragments (median of 23,727 per nucleus) and high TSS enrichment of “barcode A” (ATAC) reads of 7.3 (Fig. 2f-g), comparable to the “ATAC-only” control for SW620 T0a (Fig. 2h-i). The distribution of fragment sizes deviated from traditional ATAC profiles, likely due to the additional tagmentation step, but did not impact on fragment enrichment in accessible chromatin regions.

### Convergent evolution of the epigenome across tumour clones during 5-FU chemotherapy resistance evolution

We applied dATAC to investigate genetic and epigenetic co-evolution through chemotherapy treatment. We subjected SW620 cells to repeated cycles of 5-FU treatment at their IC50 concentration, each cycle consisting of 4 days of 5-FU exposure followed by 4-5 days of treatment holiday. Cultures were maintained in replicates and cells were harvested before treatment (T0), after one (T1) and four (T4) cycles of 5-FU treatment as well as after acquiring drug tolerance (T11/T12) (Fig. 3a). Each sample was subjected to both “ATAC-only” single-omics and dATAC multi-omics analysis to facilitate assay comparison.

**Figure 3.**
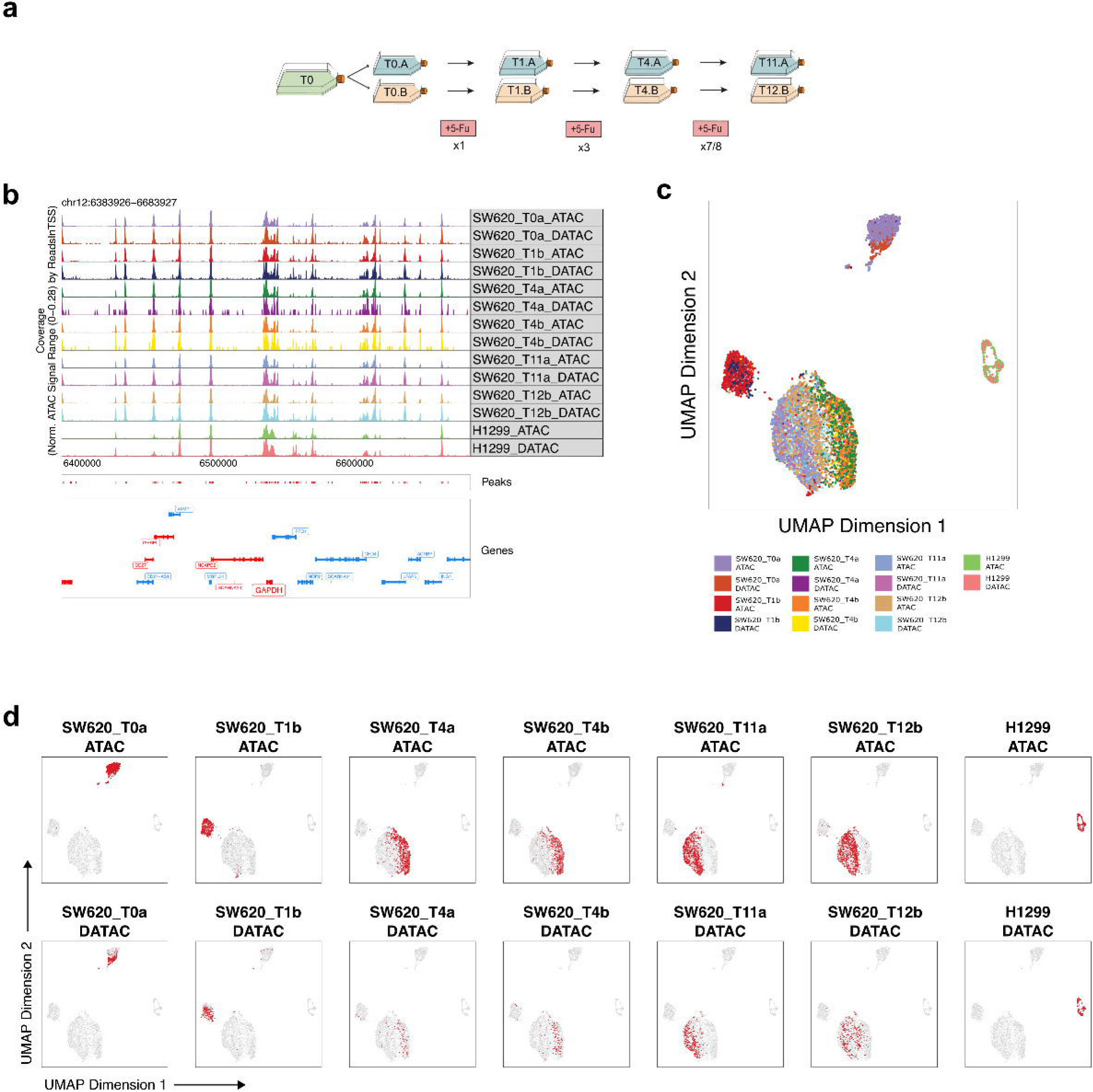
Epigenetic characterisation of SW620 cells confirms accuracy of single cell dATAC multi-omics measurements. (a) Schematic overview of treatment schedule for SW620 cells with 5-fluorouracil (5-FU). Indicated time points were subjected to both single cell “ATAC-only” and multi-omics dATAC profiling. (b) Exemplary peak accessibility profiles for the GAPDH locus show strong concordance between matched samples processed with either “ATAC-only” (ATAC) or multi-omics dATAC assay. (c) UMAP co-clustering of single-omics ATAC and matched dATAC data revealed strong clustering based on cell line (SW620 vs. H1299) and treatment exposure (SW620 only). (d) UMAP with highlights showing single cell accessibility data for individual samples from (c) revealed strong overlap in clustering for matched samples processed with either single-omics ATAC or multi-omics dATAC assay.

Exemplary peak accessibility profiles of the GAPDH locus showed strong concordance between matched samples processed with either method (Fig. 3b). UMAP co-clustering of single-omics ATAC and matched dATAC data revealed distinct cluster separation of single cells based on the cell line (SW620 vs. H1299) and treatment exposure (SW620 treatment timepoints) (Fig. 3c), and strong overlap for individual cell lines and treatment timepoints subjected to both analysis methods (Fig. 3d). Collectively, these findings illustrate the high quality of dATAC data and reveal chromatin remodelling through chemotherapy exposure.

We used genome-wide CNA data to reconstruct the phylogenetic history of 513 SW620 cells exposed to long-term chemotherapy, where dATAC had produced high quality data (Fig. 4a, b, Methods). The phylogeny was divided into eight major genetic clades with distinct karyotypes. Clades 1, 2 and 3, which largely comprised cells from the final time point (T11/T12), shared losses in chromosomes 1 and X, which were not observed in any of the other clades. Clades 4 and 5 contained most of the untreated (T0) cells and differed as clade 4 had a gain in chromosome 5, which was shared with clades 1, 2, 3 and 8, while clade 5 had a gain in chromosome 7, which was shared only with clade 6. Clades 6, 7 and 8 comprised a mixture of all time points, while clade 7 displayed the greatest genetic variability as it appeared genome doubled. Analysing clade size over time (Fig. 4c) revealed initial expansions of clades 6 and 7 after 1 and 4 treatment cycles, respectively, which then gave way to the expansion of clades 1, 2 and 3 after 11/12 cycles of treatment. This behaviour was consistent across replicates (Fig. 4c), suggesting that clades 1-3 were selected for under 5-FU treatment in a repeatable, and thus potentially predictable, way.

**Figure 4:**
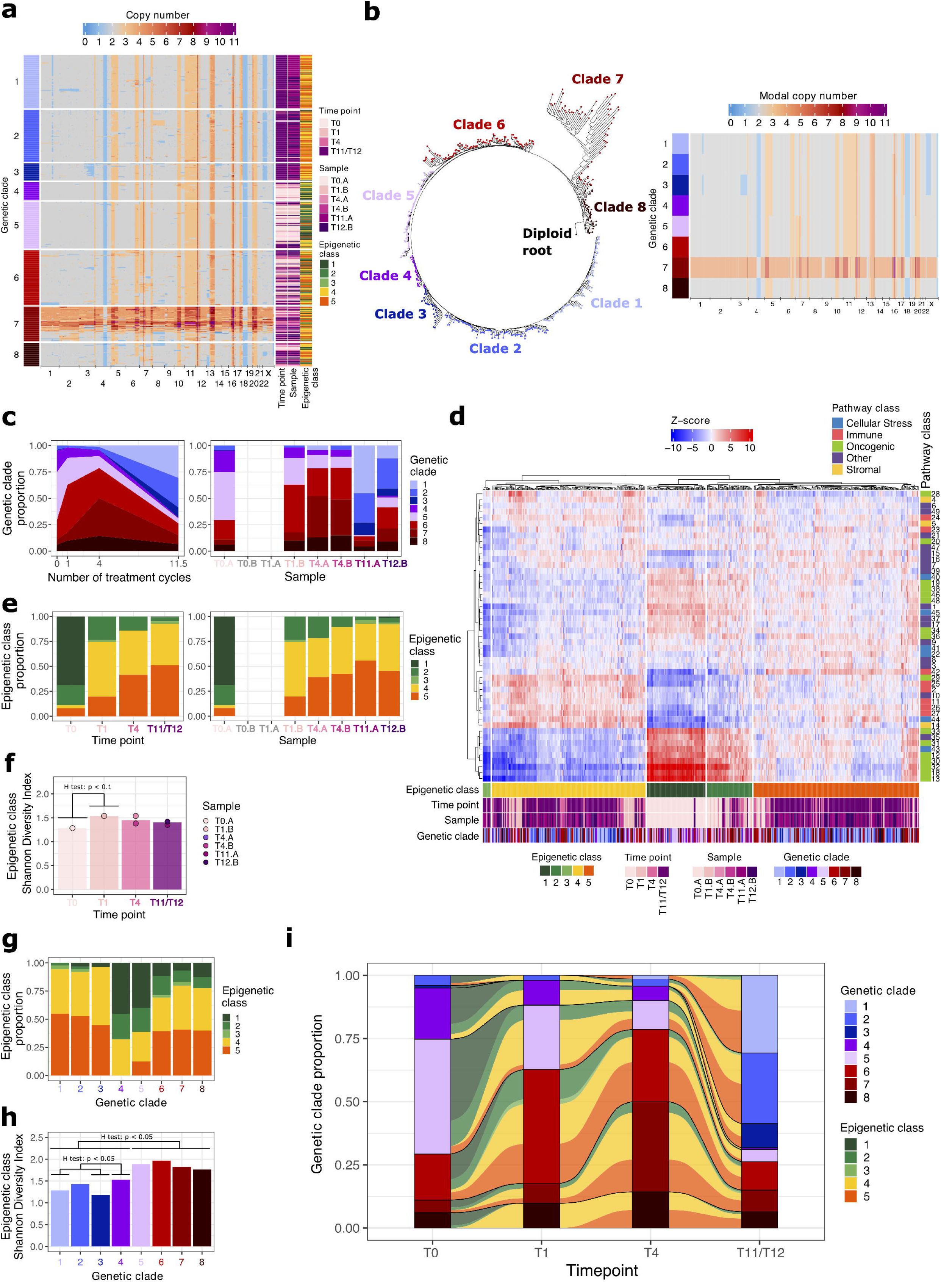
(a) CNA profiles of individual cancer cells evolved during in vitro 5-FU exposure. (b) Phylogenetic analysis divided cells into 8 major clades with distinct modal copy number (clades also depicted in a). (c) Clade sizes evolved through 5-FU exposure with clades 1, 2 and 3 likely selected by chemotherapy (left panel). Similar clade sizes were observed in a replicate evolution experiment showing reproducibility of the clonal dynamics (right panel). (d) Clustering of chromatin accessibility profiling for hallmark pathway loci divides cells into 5 epigenetic classes. Refer to Extended Data Fig. 3a for pathway names. (e) Epigenetic class sizes also evolved through 5-FU exposure (left panel) and very similar changes in epigenetic class size were observed in the replicate evolution experiment (right panel). (f) Shannon diversity of epigenetic class by timepoint shows largely stable epigenetic class diversity after long-term 5-FU exposure, with only a slight increase at T1. (g) Proportion of cells in each epigenetic class within each genetic clade. (h) Shannon diversity of epigenetic classes within each genetic clade. Clades 1-3 had lower epigenetic class diversity than clades 4-8. (i) Change in epigenetic class proportion within each genetic clade through 5-FU chemotherapy exposure. Bar clade 4, which comprised less than 5 cells from T1 onwards, all other genetic clades showed repeated (convergent) clonal evolution of the epigenome towards epigenetic classes 4 and 5 through 5-FU chemotherapy exposure.

To investigate epigenetic evolution across these genetic clades, we next used the chromatin accessibility data to predict activity scores for relevant hallmark pathways^21^ (Fig. 4d, Methods). By clustering cells according to their activity scores across all pathways, we separated them into five epigenetic classes (Fig. 4d, Methods). These classes were also detected in cells (n = 4,729) subjected to “ATAC-only” single-omics (Extended Data Fig. 2a-d, Methods). We then considered how the proportions of these epigenetic classes changed over time (Fig. 4e). Most untreated cells (T0) were in classes 1 and 2, with a few cells also in classes 4 and 5. After a single cycle of treatment, class 3 appeared, and pre-existing class 4 increased in frequency (T1), followed by further increases in frequency of classes 4 and 5 through repeated drug exposure (T4/T11/T12). The overall epigenetic class diversity in the tumour cell population was relatively stable (Fig. 4f). This suggested that epigenetic classes 4 and 5 enabled cells to adopt a phenotype that was advantageous in the presence of drug. The same pattern of epigenetic convergence towards classes 4 and 5 was consistently observed across experimental replicates (Fig. 4e), pointing towards the repeatability, and thus potential predictability, of epigenetic evolution under 5-FU treatment. In ATAC-only profiled cells, similar dynamics were observed (Extended Data Fig. 2e), with some differences in the timing of class emergence, and the overall diversity of the population (Extended Data Fig. 2f).

To infer cell phenotypes based on chromatin accessibility patterns, we considered the pathway activity scores of epigenetic classes 2-5 relative to those of class 1 (Extended Data Fig. 3a), as class 1 consisted entirely of untreated cells (bar 2 cells (out of 70) from the final time point). In classes 2-5, we observed significantly reduced accessibility of many oncogenic pathways (e.g., DNA repair, mTORC1, MYC and Notch), with only those genes typically upregulated by KRAS activation (KRAS signalling up) showing significant increase in accessibility. Cellular stress pathways related to DNA repair (unfolded protein response, UV response down) as well as pathways related to metabolism (adipogenesis, OXPHOS) also displayed significant reductions in accessibility in classes 2-5. Finally, the epithelial-mesenchymal transition pathway showed the largest increase in accessibility among all pathways. Interestingly, the epigenetic classes seem to form a continuum where, class 2 was typically more similar to class 1, while class 3, followed by class 4, were typically more distinct from class 1, and class 5 was an intermediate state (Extended Data Fig. 3a,b). Given the order in which the different epigenetic classes appeared and/or expanded in the population (Fig. 4e, Extended Data Fig. 3b-e), this suggested that treatment concurrently drove the epigenetic composition of the tumour cell population in two directions: there was both a large initial shift from class 1 to classes 3 and 4 and a more gradual transition from class 1 to class 2. Class 5 likely represented cells that evolved towards class 1 from classes 3 and 4, or away from class 1 from class 2. To further explore differences between the epigenetic classes, we predicted transcription factor (TF) family scores from the chromatin accessibility data (Methods, Extended Data Fig. 2f-j), observing significant enrichment of FOS and JUN-related factors in epigenetic classes 4 and 5 compared to untreated cells, consistent with previous studies associating FOS to 5-FU resistance in CRC^22^.

Using the multi-omic readout of dATAC, we then combined genomic and epigenomic data to measure the clonal evolutionary history of epigenome alteration through treatment. The genetic clades which increased in frequency through treatment, i.e. which were selected by drug (clades 1,2 and 3), were predominantly composed of epigenetic classes 4 and 5 (Fig. 4g) and had lower epigenetic diversity compared to the other genetic clades (Fig. 4h). Epigenetic classes 4 and 5 were also consistently selected for over time across all genetic clades that remained sizeable after treatment (Fig. 4e,i). Further, epigenome evolution was repeatable within clades (Fig. 4i), mirroring our earlier observations of repeated genome (clade) and epigenome (class) evolution across experimental replicates (Fig. 4c,e). This was highly suggestive that epigenetic rewiring occurred independently of underlying genome alteration. Taken together, these results suggest that continued exposure to 5-FU drives natural selection of pre-existing genetic and epigenetic variation, selecting for tumour cells that have both a particular genome and an advantageous epigenome. There is likely then further epigenome remodelling through treatment. These data support the idea that genome and epigenome co-evolution are predictable.

### Chromosomal alterations are associated with reorganisation of chromatin architecture

We leveraged the co-measurement of copy number and chromatin accessibility profiles in single cells obtained with dATAC to study the interrelationship between CNAs and epigenetic remodelling of the altered genomic regions. Using the CNA profiles of individual tumour cells undergoing 5-FU treatment (Fig. 4a), we identified three subclonal copy number alterations at high frequency in the cell population (Fig. 5a, Extended Data Fig. 4a, methods): chromosome 5 gain, and losses in chromosome 1 and X, respectively. For each of these copy number alterations, we separated cells into two clones (one with and one without the alteration of interest), generated pseudo-bulk ATAC signal for each clone and compared the ATAC signal in peaks between the clones (Fig. 5b-d, Methods).

**Figure 5:**
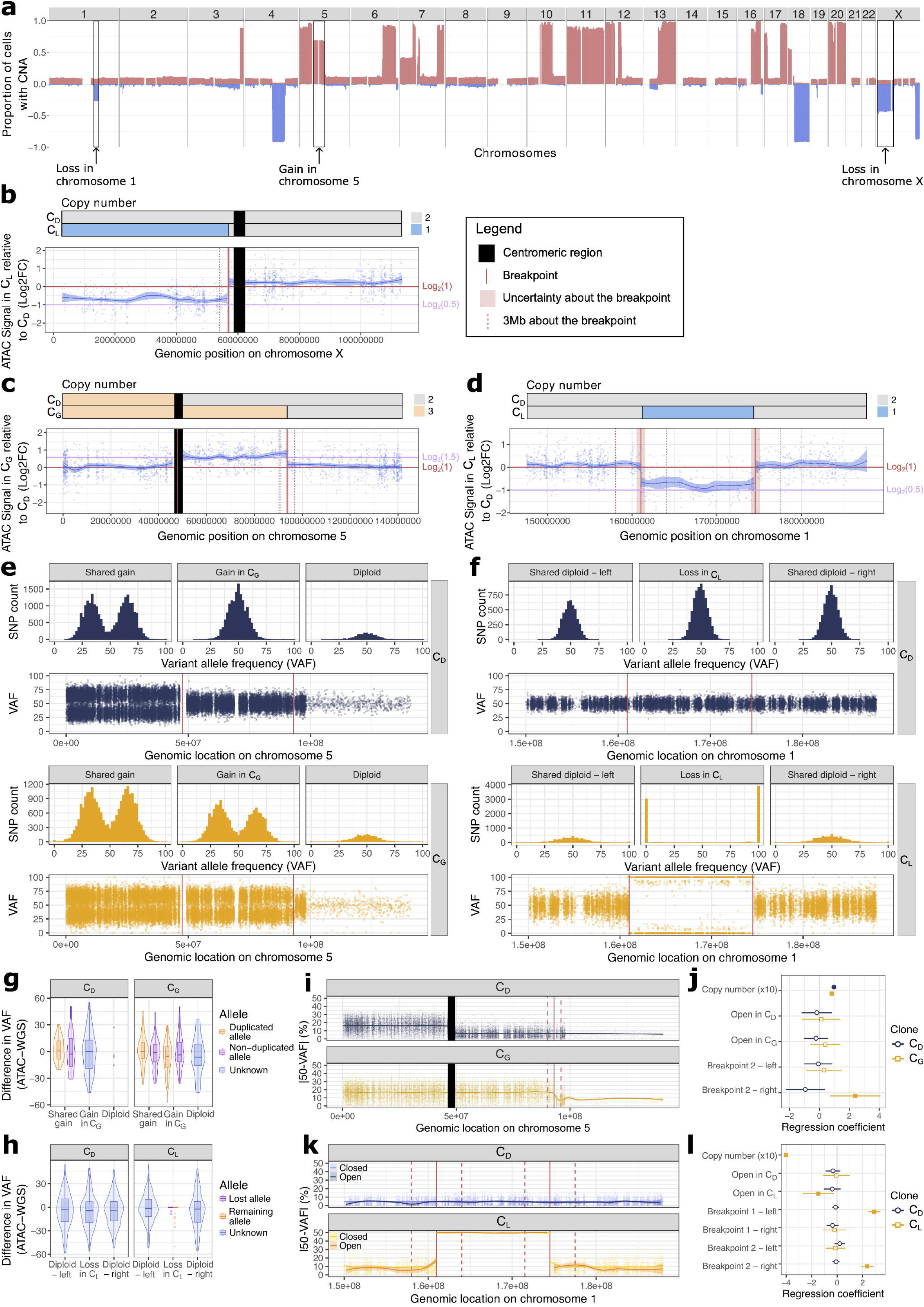
(a) Genome-wide frequency plot of CNAs in the cancer cell population (from Fig. 4a,d) highlighting three high-frequency subclonal copy number alterations: a gain in chromosome 5 and losses in chromosome 1 and X, respectively. (b-d) Comparison of the ATAC signal in peaks between clones comprising cells with the copy number alteration of interest (C_G_ for the gain in chromosome 5, C_L_ for the losses in chromosomes 1 and X) and clones comprising cells that do not have the copy number alteration of interest (C_D_). The three CNAs were associated with differences in chromatin accessibility proportional to the differences in allelic copy number, indicating a stochiometric difference in accessibility. (e-f) The VAFs of heterozygous SNPs in the regions of interest on chromosomes 5 and 1, computed from the WGS component of the dATAC data, followed expected distributions in clones with (C_G_, C_L_) and without (C_D_) CNAs. (g-h) Difference in VAFs computed using the ATAC and WGS components of the dATAC data for heterozygous SNPs on the chromosome 5 and 1 regions of interest in clones with (C_G_, C_L_) and without (C_D_) CNAs. All distributions were consistent with equal allele accessibility. (i) For clones C_G_ and C_D_, comparison of smoothed absolute distance of VAFs from 50% (computed from the WGS component of the dATAC data) for heterozygous SNPs on the chromosome 5 region of interest. (j) Coefficient plot for the linear regressions predicting |VAF-50| in clones C_G_ and C_D_ based on copy number, proximity to breakpoints and presence in open vs closed chromatin regions. Shaded shapes highlight significant (p < 0.05) predictor variables. Significantly higher than expected values of |VAF-50| were observed in clone C_G_ for SNPs in the common diploid region flanking the second breakpoint. (k) For clones C_L_ and C_D_, comparison of smoothed absolute distance of VAFs from 50% (computed from the WGS component of the dATAC data) for heterozygous SNPs on the chromosome 1 region of interest. (l) Coefficient plot for the linear regressions predicting |VAF-50|in clones C_L_ and C_D_ based on copy number, proximity to breakpoints and presence in open vs closed chromatin regions. Shaded shapes highlight significant (p < 0.05) predictor variables. Significantly higher than expected values of |VAF-50| were observed in clone C_L_ for SNPs in the common diploid regions flanking both breakpoints. Values of |VAF-50| were significantly lower for SNPs in open chromatin regions compared to SNPs in closed chromatin regions.

For all three CNAs, we found that, at the length scale of the CNAs, the differences in chromatin accessibility between the clones with the CNA and without the CNA were proportional to the differences in allelic copy number due to the alterations (Fig. 5b-d). Under the common assumption of equal allele accessibility, this suggests the absence of a mechanism to remodel chromatin accessibility in response to somatic chromosomal aberrations. We further sought to determine whether there were any significant alterations in chromatin accessibility in regions flanking the breakpoints, i.e., at shorter length scales. We sampled 1,000 random loci from the genomic segments to the left and right, respectively, of each breakpoint. We then compared the mean log2 fold change (L2FC) in ATAC signal (between CNA clones) within 100kb-1.5Mb to the left (or right) of each breakpoint to the distributions of equivalent mean L2FCs in ATAC signal obtained using the random loci (methods). While, for chromosome 1, the chromatin accessibility in clone C_L_ (cells with chr1 loss) was significantly lower than expected in the diploid region within 100kb of the left breakpoint (ratio of true value to null mean: -4.05, FDR-adjusted p = 0.096, Extended Data Fig. 4b), there was no other evidence of chromatin remodelling near breakpoints (Extended Data Fig. 4b,c). This indicated remarkable stoichiometry of chromatin accessibility at shorter length scales near CNA breakpoints. We note, however, that the sparsity of the chromatin accessibility data in small genomic regions limited the power to detect accessibility changes around breakpoints, if such changes exist.

We used single nucleotide polymorphisms (SNPs) to examine allele-specific chromatin accessibility. Exploiting the high library complexity of dATAC-profiled cells, we called heterozygous SNPs in the genomic regions of interest in chromosome 1 and 5 (Methods). The loss in chromosome X was excluded, as it represented a “somatic” reversal of a duplication event (SW620 cells are male with a somatic duplication of chromosome X), and therefore all SNPs in this region were homozygous. For the previously defined clones with and without CNAs, we examined SNP variant allele frequencies (VAFs) separately using the WGS (Fig. 5e,f, Methods) and the ATAC (Extended Data Fig. 4d,e, Methods) components of the dATAC data. The WGS-derived VAFs (Fig. 5e,f) fully recapitulated expected VAF distributions, with unimodal distributions centred around 50% for the diploid regions, and bimodal distributions centred around 33% and 66% for the gain regions and around 0% and 100% in the loss region. The ATAC-derived VAFs (Extended Data Fig. 4d,e) showed no evidence of allele specific differences in accessibility, but we note the higher noise due to read count sparsity. To further test for allele-specific differences in chromatin accessibility, we compared the SNP VAFs computed using the ATAC and WGS data (Fig. 5g,h, Extended Data Fig. 4f,g). In diploid regions, the distributions of the differences in ATAC-derived and WGS-derived VAFs were all unimodal and approximately centred at 0. In gain regions, there was no significant difference in the distributions of the differences in ATAC-derived and WGS-derived VAFs for the duplicated and non-duplicated alleles (Wilxocon test, p > 0.05 in all cases). These findings all support the assumption of equal allele accessibility.

Finally, we examined allele specific information close to CNA breakpoints in WGS data. In clone C_G_ (gain in chromosome 5), in the diploid region close to the breakpoint we found deviations of SNP-VAFs from 50% within 1Mb of the left breakpoint (linear regression coefficient of 2.41%, p = 5.28 x 10^-3^) (Fig. 5i,j, Methods). Similarly, for clone C_L_ (loss in chromosome 1) there were deviations of SNP-VAFs from 50% within 1Mb of the breakpoints (linear regression coefficients of 2.89% (p < 10^-16^) and 2.47% (p < 10^-16^), respectively) (Fig. 5k,l, Methods). dATAC WGS reads (B barcodes) contain only a minority of open chromatin reads (< 1%), and therefore the SNP-VAF deviations around breakpoints were most likely suggestive of complex aberration processes that resulted in “jagged” breakpoints, rather than alterations in chromatin accessibility near breakpoints.

Overall, our results demonstrate how multimodal measurement by dATAC provides the means to investigate the association between copy number alterations and modifications in local chromatin architecture.

### Mapping epigenetic states to cell genotypes in single cells from patient-derived metastatic tissue

We applied dATAC to fresh-frozen surgical biopsy material from two patients (P05 & P06) with metastatic peritoneal disease derived from primary colorectal cancer and also performed single-omic ATAC profiling on the same samples (Fig. 6a). Patient material required a modified protocol for nuclei extraction (see methods).

**Figure 6.**
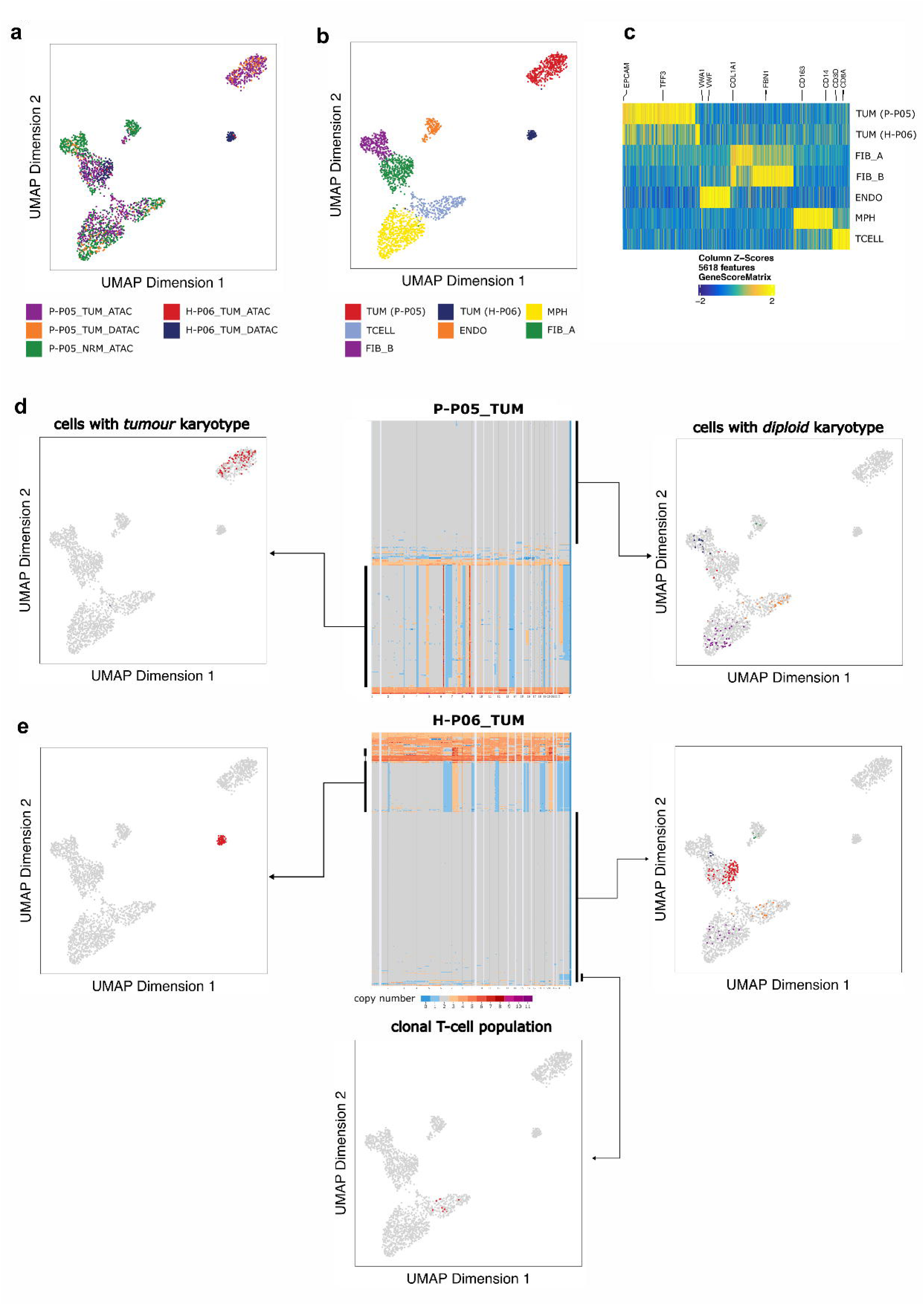
Tissue samples were collected from patients with oligometastatic peritoneal disease derived from colorectal cancer. (a) UMAP representation of chromatin accessibility data for one peritoneal metastasis and one adjacent normal tissue biopsy from patient P05 and one peritoneal metastatic tissue biopsy of patient P06. Both metastatic samples were subjected to matched single-omics “ATAC-only” and multi-omics dATAC assay analysis. (b) UMAP data from (a) coloured according to Seurat clustering identified distinct tumour and stromal cell populations in peritoneal tissue. TUM - tumour epithelial cells, MPH - macrophages, ENDO - endothelial cells, TCELL – T cells, FIB_A/FIB_B - fibroblast cell populations. (c) Heatmap analysis depicting gene score z-scores for each identified cluster revealed cell type specific gene accessibility patterns in peritoneal tissues. Significantly enriched cell-specific differentiation markers are highlighted. (d-e) Integrated analysis of paired chromatin accessibility and WGS data from single cells in peritoneal metastatic tissue from patient P05 (d) and P06 (e). Cross-referencing of both data modalities showed that cells with mainly diploid cell karyotypes exclusively map to stromal cell compartments, whereas cells with tumour karyotypes exclusively map to epithelial tumour cell populations in the gene accessibility UMAP. A population of T cells in P06 were identified as having a clonal loss of chromosome X (e), with an abundance of 35% (8/23) in the total T-cell population.

Clustering of chromatin accessibility data from all three biopsies identified tumour and stromal cell types with distinct epigenetic profiles (Fig. 6b-c). Analysis of the WGS component of the data revealed presumed tumour cells carrying a large burden of CNAs and detected subclones with distinct CNAs in the population. Most sequenced cells in each sample were diploid, and were presumably stromal cells that did not carry the genomic alterations of the tumour cells (Fig. 6d-e). To evaluate this possibility, we used the paired chromatin accessibility data to identify the cell type of each of cell. As expected, the aneuploid cells were cancer cells, and the diploid and near-diploid cells were exclusively stromal (Fig. 6d-e). Of note, dATAC detected somatic alteration and clonal expansion of T cells: 8/23 (35%) of T cells detected in patient P06 harboured a clonal loss of chromosome X (Fig. 6e, bottom panel).

## Discussion

Here we have presented dATAC, a technology for simultaneous assessment of genomic and epigenomic profiles in single cells, that produces multi-modal data of the same quality as leading single-omics approaches. The protocol requires minimal specialist equipment, is carried out in a convenient one-pot reaction format and is suitable for use on cell lines and fresh-frozen patient materials.

The simultaneous profiling of the genome and epigenome in individual cells by dATAC enables the study of the somatic evolution of the epigenome. Somatic genetic alterations record the clonal ancestry of cells: overlaying epigenetic information on top of a phylogenetic tree reveals the evolutionary history of epigenetic heterogeneity. Here, we have used this approach to delineate how chemotherapy initially selects for genetic clones in colorectal cancer, likely because of the epigenetic makeup of the clones, which then undergo convergent and predictable epigenetic rewiring through continued treatment exposure. We also mapped the epigenetic consequence of genetic alteration, revealing no evidence of epigenetic buffering as a consequence of copy number alteration. These are examples of how multimodal profiling is a window to explore the fundamental interlinking of somatic epigenome alteration, genetic mutation and clonal evolution.

We note that the dATAC approach of sequential rounds of tagmentation with distinct tagment barcodes could in theory be extended to more than two rounds of tagmentation, enabling multiplexed measurement of further genetic or epigenetic marks (e.g. CUT&Tag approaches for histone marks^23, 24^).

In summary, dATAC is an extensible and “open source” methodology for high-quality multi-omic profiling in single cells, that offers a powerful window into the interlinked somatic evolution of the genome and epigenome.

## Contributions

Study conception and design: TG

dATAC method design and implementation: MM

Experimental work: MM and ASt

Experimental evolution conception and design: FW and TG

Data analysis conception and design: MM, CC and TG

Bioinformatics analysis: CC, MM, QG, HG, FW

Provision of patient materials: PK, DDB and JM

Critical analysis of data: ASt, AMB and Aso

Lead manuscript writing: MM, CC and TG

## Acknowledgements

This study was principally funded by Cancer Research UK by programme grant DRCNPG-May21_100001 to TG. We thank the ICR’s Genomics Facility for their sequencing support as well as Erica Oliveira, Julia Rezende da Silva and Caitlin Davies for their help with tissue culture.

## Methods

### Preparation of cell lines

MCF7 cells were cultured in Dulbecco’s Modified Eagle Medium (DMEM) supplemented with 10% fetal bovine serum (FBS), 50 μg/mL streptomycin sulfate, 100 units/mL penicillin, 2.5 mmol/L L-glutamine (1% (v/v) PSG, Sigma) and 10^−8^ mol/L 17-ß-estradiol (E2, Sigma-Aldrich) and were incubated at 37°C with 5% CO_2_ in a humidified atmosphere. SW620 and H1299 cells were cultured in DMEM (high glucose, pyruvate) medium (Gibco) supplemented with 10% FBS, 100 μg/mL streptomycin sulfate, 100 units/mL penicillin (Gibco) and incubated at same conditions as MCF7 cells. For treatment with 5-FU, SW620 cells were treated with 22.97 μM 5-FU (IC50 value) in standard medium for 5 days. Cells were harvested at the end of the 5 day treatment and immediately subjected to ATAC and dATAC preparations.

### Nuclei preparation from cell lines

Trypsinised cells from cell lines were washed in PBS/0.04% BSA and quantified using LUNA FL cell counter (Logos Biosystems). Aliquots of 50,000 cells were washed again with PBS/0.04% BSA and spun down at 300g for 5 min. Cell pellets were resuspended in 50ul lysis buffer containing 48.5ul RSB (10mM Tris-HCl pH 7.4, 10mM NaCl, 3mM MgCl_2_ in ddH2O), 0.5ul 10% NP-40, 0.5ul 10% Tween-20 and 0.5ul 1% Digitonin and incubated on ice for 3 min. Subsequently, lysed nuclei were diluted with 1ml of ice-cold RSB-T buffer (RSB + 0.1% Tween-20 (v/v)), inverted 3 times and spun down at 500g for 10 min at 4°C in a swing-bucket rotor centrifuge. All remaining supernatant was carefully removed before proceeding with the tagmentation step.

### Nuclei preparation from human patient tissue

Frozen tissue biopsies were transferred onto a pre-cooled petri dish on dry ice and a ∼1mm^3^ piece was dissected using a pre-cooled surgical blade. After filling a pre-cooled 2ml Dounce homogenizer on ice with 700ul nuclei extraction buffer (10mM Tris-HCl pH 7.4, 10mM NaCl, 3mM MgCl_2_, 0.1% Tween-20 (v/v), 0.1% NP-40, 1mM dithiothreitol and 2% bovine serum albumin (BSA) in H2O) the dissected frozen tissue was transferred into the homogenizer and the tissue allowed to sink. Immediately after that, 10 gentle strokes were performed with pre-cooled Dounce pestle “A” and the homogenized tissue was incubated on ice for 5 min. Subsequently, the homogenate was filtered through a 100uM mesh into a 1.5ml tube. The homogenizer was rinsed with 300ull ice-cold nuclei extraction buffer, which was filtered and added to the homogenate, and the combined nuclei extract was spun down at 500g for 10min at 4°C in a swing-bucket rotor centrifuge. All remaining supernatant was carefully removed, and nuclei were resuspended in 500ul PBS/2% BSA. Luna FL cell counter analysis was performed to confirm successful nuclei extraction and quantify nuclei counts. 16,000 nuclei were then aliquoted, further diluted with 1ml ice-cold PBS/2% BSA and spun down at 500g for 10 min at 4°C, after which all remaining supernatant was carefully removed before proceeding with the tagmentation step.

### Barcode adapter loading on custom Tn5

Diagenode Tn5 unloaded enzyme was purchased and loaded with double stranded adapter following the manufacturer’s recommended protocol. For adapter hybridisation barcoded iTSM and ME-REV oligonucleotides were used.

### Open chromatin tagmentation of nuclei

Nuclei pellets from either cell lines or tissue were carefully resuspended in 50ul tagmentation mix (25ul 2x TD buffer (20mM Tris-HCl pH 7.6, 10mM MgCl_2_, 20% dimethylformamide (v/v) in H2O), 16ul PBS, 0.5ul 10% Tween-20, 0.5ul 1% Digitonin, 6ul 10% BSA and either 2ul custom barcoded Tn5 (cell lines) or 1ul custom barcoded Tn5 and 1ul H2O (tissue)). Nuclei tagmentation mixes were incubated at 37°C for 30min on an orbital shaker at 800rpm. Tagmentation reactions were stopped by mixing the reaction with 50ul 2x STOP buffer (10mM Tris-HCl pH 8, 20mM EDTA and 2% BSA in H2O). A further 800ul ice-cold RSB-T were added and nuclei spun down at 500g for 10 min at 4°C. Nuclei pellets were resuspended in 50ul sorting buffer (1x STOP buffer, 100ng/ml DAPI) and kept on ice until proceeding to plate sorting.

### Preparation of lysis plates

384 unique i5 indexing primers (IDT, supplemental table S1) were ordered in 384 plate format and diluted to 8uM concentration in a separate primer stock plate. 384-well sorting plates with primers were prepared in batches by multi-dispensing 1ul of the primer stock plate. After spinning down sorting plates, all liquid volume was allowed to evaporate and plates with lyophilised primers were sealed and stored at -20°C until further use. Immediately before sorting, one primer plate per sort was thawed and 1ul of lysis buffer was dispensed per well (ATAC plate: 0.5ul 2x TD buffer, 0.2ul Qiagen Protease, 0.03ul 10uM A14-LNA-ME, 0.1ul 1% SDS, 0.17ul H2O; dATAC plate: 0.5ul 2x TD buffer, 0.2ul Qiagen Protease, 0.04ul 10uM A14-LNA-ME, 0.26ul H2O). Plates were then spun down and kept on ice until being used for sorting.

### Single nuclei sorting using CellenOne system

Nuclei resuspended in sorting buffer were filtered through 20um cell strainer immediately prior to sorting on the real-time image-based CellenOne F1.4 sorting system (Scienion). Ambient instrument conditions were set to 55% humidity and the temperature of the 384-plate holder was set to -1°C below dew point temperature. 5ul of nuclei suspension were taken up with a PDC-M nozzle and 100 nuclei were assessed for adjustment of size, elongation and DAPI intensity settings. DAPI-positive single nuclei were then sorted into individual wells of a lysis plate and visually confirmed after sorting to exclude wells with multiple nuclei. For ATAC/single-modal NGS preparations, single nuclei from different bulk open chromatin tagmentation, each with a unique Tn5 inline barcode, were sorted into a single well to maximise cell throughput per library preparation, usually with 6 nuclei per well. After successful sorting, plates were spun down at 3,000g for 5 min at 4°C and stored at -20°C until library preparation.

### Library preparation protocol – ATAC-seq (single-modal NGS)

Please note that every step in the single- and multi-modal library preparation protocol requires spin down using a bench-top centrifuge, thorough vortex-mixing followed by another spin down of the plate after every addition of reagents indicated below, which is critical to ensure optimal protocol performance. An overview of indexing primers and the template switching oligo can be found in supplemental Table S1.

Frozen sorted plates were thawed and spun down at 3,000g for 2 min at 4°C, and sorted nuclei were then lysed in a 384-well Biorad C1000 touch thermocycler at 55°C for 15 min, 70°C for 10 min and storage at 10°C. After adding 1ul 1% Triton-X 100 to quench SDS present in the lysate, 1ul of NEB Q5 2x Master Mix was added. Gap-filling of tagmented DNA and introduction of the Nextera-style i5 handle, using the switching oligo (A14-LNA-ME) as a template, was carried out using following cycling program: Gap-filling at 72°C for 5 min followed by 10 cycles at 98°C for 30s, 59°C for 20s and 65°C for 60s, with final storage at 10°C. The gap-filled and extended fragments were then PCR-amplified by adding the following PCR mix to the 3ul of sample volume: 3ul NEB Q5U 2x Master Mix, 0.04ul 200uM TruSeq i7 plate index primer, 1.96ul H2O. For plates with 6 nuclei per well PCR amplification was carried out using following conditions: 98°C for 30s, 12 cycles at 98°C for 30s and 65°C for 75s, with final elongation at 72°C for 2min and storage at 10°C. All wells were pooled and purified with the Zymo DCC-5 kit after diluting the pool 5-fold with DNA binding buffer. Final ATAC libraries were eluted in 40ul and quantified using Agilent Tapestation HSD5000 screentape on an Agilent Tapestation 4200 system.

### Library preparation protocol – dATAC-seq (multi-modal NGS)

Frozen sorted plates were thawed and spun down at 3,000g for 2 min at 4°C, followed by >24h incubation at +4°C to ensure full penetration of nuclei with lysis buffer. The plate was spun again, and sorted nuclei were then lysed in a 384-well Biorad C1000 touch thermocycler at 50°C for 1h, 70°C for 10 min and subsequently stored at 10°C. This step ensures the release of both pre-tagmented open chromatin-derived fragments as well as previously inaccessible genomic DNA. After lysis of the single nuclei, tagmentation of now freely accessible whole genomic DNA was carried out. While on ice, 1ul of Tn5 mix, consisting of 0.5ul 2x TD buffer, 1.75*10^-3^ul barcoded Tn5 (different from the Tn5 barcode used for the ATAC preparation before sorting) and 0.498ul H2O, was added to each well and the plate was incubated at 55°C for 10min, followed by storage at 10°C. The plate was again placed on ice and the tagmentation reaction was stopped by adding 1ul neutralisation mix (0.5ul Qiagen Protease, 0.01ul 10% Tween-20 and 0.49ul H2O) and incubated at 50°C for 15min, followed by 70°C for 10 min and storage at 10°C. Gap-filling and i5-handle switching were carried out as described for the ATAC-Seq preparation above. Open-chromatin and closed chromatin-derived fragments were subsequently co-amplified in the same well using 5ul PCR mix (3.5ul NEB Q5U 2x Master Mix, 0.04ul 200uM TruSeq i7 plate index primer, 1.46ul H2O) with the following cycling conditions: 98°C for 30s, 9 cycles at 98°C for 30s and 65°C for 75s, with final elongation at 72°C for 2min and storage at 10°C. 4.5ul (half volume) from each well were pooled and purified and quality controlled as described for ATAC-seq libraries. The plate with remaining reaction volumes was stored at -20°C. For deep-sequencing of individual single cell libraries, the plate was returned to the PCR cycler for additional 8 cycles of PCR. Select single cell libraries were then individually purified using 2 volumes of cleanNGS beads (CleanNA) and quantified using Agilent Tapestation HSD5000 screentape analysis.

### Library pooling and sequencing

ATAC and/or dATAC NGS libraries were pooled and subjected to residual primer digestion using thermolabile exonuclease-1 (NEB), followed by 2x (v/v) cleanNGS bead purification. All library pools were then sequenced on the Novaseq6000 platform on 300 cycle S4 flow cells with run mode 160-8-10-160 and 15% PhiX spike-in.

### Chromatin accessibility analysis

After sequencing the paired reads data was first demultiplexed based on i5 (wells) and i7 (plate) indexes. To further separate the raw data from each well into individual nuclei (ATAC/single-modal prep) or open vs closed chromatin from a single nucleus (dATAC/multi-modal prep), demultiplexed fastq containers were processed with ultraplex v1.2.5 software by providing a list of all utilised Tn5 inline barcodes and the following parameters: *-q 0, --fiveprimemismatches 1, --adapter2 CTGTCTCTTATACACATCTGACGCTGCCGACGA --adapter CTGTCTCTTATACACATCT*. Because the first 8bp of fastq read2 represent the Tn5 inline barcode, fastq read2 was used as “read1” input for ultraplex to utilise its barcode detection function. The 20bp universal adapter sequence that follows the Tn5 inline barcode was removed, and all reads trimmed to a length of 35bp using seqtk function trimfq. After preprocessing the data into individual fastq containers for individual nuclei, paired reads for the barcodes representing first round tagmentation (open chromatin) were mapped to GRCh38 (Gencode p13_v41) using bowtie2 (v2.4.2) with the following parameters: *-p 4 --local --very-sensitive -X 1000 -k 5 --no-mixed --no-discordant*. Unique cell barcodes were introduced into each bam file via CB tag with a custom script using samtools and awk. Subsequently, samtools software was used to sort and index bam files and consensus reads were generated from PCR duplicates using the gencore tool. All bam files (representing ATAC data of individual nuclei) from the same biological sample were then merged into a single bam file using samtools’ merge function.

Merged sample-level bam files were then loaded into ArchR software and processed following the recommended guidelines to generate UMAP clustermaps, genomic region peak profiles as well as QC assessments. TSS enrichment >=4 and unique fragments >=5000 were used as cut-offs for subsequent analyses, whereas ATAC-only and DATAC-derived open chromatin data were integrated into the same analysis using the harmony tool.

### Whole genome profile analysis

Sequencing data from single cells undergoing the DLP+ or dATAC protocol was demultiplexed into dual-index fastq files and used as input for the workflow automation pipeline designed for the DLP+ protocol (https://github.com/shahcompbio/single_cell_pipeline) with default settings. First, raw reads were adapter-trimmed using TrimGalore, mapped with BWA aln to the hg19 reference genome and deduplicated using picard MarkDuplicates. Subsequently, copy number calling was performed using the HMMcopy tool with reads segmented into non-overlapping 500kb genomic regions.

### Construction of the single-cell phylogeny from copy number data obtained via dATAC

#### Additional filtering of the copy number data

To reconstruct a single-cell phylogeny across all time points and samples, the copy number data obtained from HMMCOPY was further filtered to exclude low-quality cells and low-mappability bins, which correspond to regions of the genome that are difficult to sequence. Of the cells that passed the ATAC QC performed in ArchR (with minTSS = 4 and minFrags = 5,000, see Chromatin Accessibility Analysis), those cells with a quality score, as defined in Laks et al^20^, above 0.75 and an S-phase probability above 0.5 were initially retained. The copy number bins were then also filtered to only retain bins with a mappability score above 0.99. HMMCOPY’s S-phase classifier may miss cells that are in the early or late stages of replication and are characterised by focal alterations scattered across the genome^25^. Since such scattered patterns are not representative of true evolutionary history, but may (erroneously) bias the phylogenetic inference, remaining cells were ranked by their total number of copy number changes between consecutive bins, and the lower 90^th^ percentile of cells were retained. After these filtering steps, the copy number data included copy number calls for 513/1920 cells, or 26.7% of cells, across 4363/6206 genomic bins, or 70.3%, of the genome.

#### Creation of a minimum consistent segmentation across all cells

For each cell, a cell-specific minimum segmentation was created by merging consecutive bins (on the same chromosome) with equal copy number. A minimum segmentation across all cells was also created by merging consecutive bins (on the same chromosome) with cell-specific equal copy number (i.e., different cells may have different copy number in the merged bins). Then, the procedure introduced in Watkins et al^26^ was used to generate a minimum consistent segmentation across all cells. In more detail, a maximal set of breakpoints was created by merging cell-specific minimum segmentations together, retaining all breakpoints and keeping track of the cell-of-origin. Then, breakpoints from different cells that were separated by a distance less than or equal to 2Mb were iteratively merged. This corrects for cell-to-cell variation in breakpoint location which can arise as an artifact from the copy number calling even when the breakpoint location is the same across cells. Internal functions from Refphase (http://bitbucket.org/schwarzlab/refphase) were used to create the minimum consistent segmentation as described above.

#### Phylogenetic reconstruction using DICE

After adding a normal, diploid cell to the final copy number data, a maximum parsimony tree was reconstructed using DICE (https://github.com/samsonweiner/DICE)^27^. Default command line options were used, except for the use of the total copy number flag -t.

#### Definition of the genetic clades

A maximal set of 104 clades was generated using mPTP v0.2.5^28^, a tool developed for species delimitation from a phylogenetic tree, with the options ‘--mcmc 50000000 --multi --mcmc_sample 1000000 --mcmc_burnin 100000 --outgroup Diploid’. These 104 clades were then iteratively merged to obtain 8 clades, by grouping neighbouring clades on the phylogeny that have similar copy number profiles.

#### Estimation of pathway expression from the chromatin accessibility data obtained via dATAC and ATAC-only

When an ArchR project is created, gene scores, which are a prediction of gene expression based on the accessibility of regulatory elements near the gene, are computed by default. These gene scores are stored in a sparse matrix, GeneScoreMatrix, which was obtained via the function getMatrixFromProject. Separate gene score matrices for the cells subjected to dATAC and ATAC-only were obtained by filtering the GeneScoreMatrix to contain only gene scores of genes involved in the pathways of interest, for (1) tumour cells subjected to dATAC included in the phylogenetic tree and (2) all tumour cells subjected to ATAC-only.

Since the sparsity of single cell ATAC data can obscure the true signal in the data, MAGIC (https://github.com/KrishnaswamyLab/MAGIC), an algorithm developed to denoise high-dimensional data, was applied to both gene score matrices to impute missing gene scores and smooth the gene scores across cells. The R implementation of MAGIC (Rmagic R package v2.0.3) was used and the algorithm was run with the function ‘magic’ using default parameters, except for the smoothing parameter, which was set to t=2. Finally, gene set variation analysis (GSVA) was performed on both imputed gene score matrices using the GSVA R package v1.50.0, with the z-score method^30^, to estimate pathway activity across cells using their z-scores.

#### Definition of epigenetic classes from the dATAC data

A Euclidean distance matrix of the pathway activity z-scores was calculated using the ‘dist’ function from the stats R package v4.3.2. The ‘hclust’ function from the stats R package was then used to perform hierarchical clustering with complete linkage. Five epigenetic classes were finally found using the ‘cutree’ function from the stats R package.

#### Assigning dATAC-derived epigenetic classes to tumour cells subjected to ATAC-only

A k-nearest neighbours (KNN) classification algorithm was used to predict the epigenetic class of tumour cells subjected to ATAC-only based on their hallmark pathway scores. The KNN classification model was trained on the hallmark pathway scores and corresponding epigenetic classes for tumour cells subject to dATAC using the ‘knn3’ function of the caret R package v6.0.94, with k = 3 neighbours. The number of neighbours was selected based on results from repeated cross-validation (10 repeats of 10-fold cross validation), which was performed using the ‘train’ function of the caret R package. We tested 2-30 neighbours and found that 3 neighbours yielded the highest accuracy of 89.3%. Epigenetic class prediction for tumour cells subjected to ATAC-only was then performed using the ‘predict’ function of the stats R package v4.3.2.

The epigenetic class classification obtained for cells in the ATAC-only dataset was validated by comparing mean hallmark pathway scores in each epigenetic class for the dATAC and ATAC-only datasets. In particular, Pearson correlation coefficients, and their respective p-values, of the ATAC-only and dATAC mean hallmark pathway scores were calculated for each epigenetic class using the ‘cor.test’ function of the stats R package.

#### Quantification of the epigenetic diversity per time point, sample and genetic clade

The Shannon diversity index for each time point, sample and genetic clade was computed using the ‘diversity’ function of the vegan R package v2.6.8, setting the index as “shannon” and a logarithm base of 2. The significance of the difference between the Shannon diversity indices of two groups was tested using the Hutcheson t-test. The multiple_Hutcheson_t_test function from the ecolTest R package v0.0.1 was used to compute the p-values of the Hutcheson t-test in a pairwise way between time points, samples and genetic clades.

#### Estimation of transcription factor (TF) family activity from the chromatin accessibility data obtained via dATAC

To study TF motif enrichments, we first added motif annotations for a motif set curated from CIS-BP^31^ to our ArchR project using the function addMotifAnnotations with motifSet set to “cisbp”. ArchR predicts single-cell TF enrichments using an extension of ChromVar^32^, which allows its application to larger datasets. ChromVar computes bias-corrected deviation scores, which measure how much the accessibility of a given TF in each cell deviates from the expected accessibility based on the average over all cells, and their z-scores. The bias corrections are performed by defining, for each peak containing a TF of interest, a set of background peaks, which have similar GC-content and average accessibility, and using these background peaks’ deviation scores to normalise the peak’s (and TF’s) deviation score. We, thus, identified background peaks using the function addBgdPeaks and computed deviation scores using the function addDeviationsMatrix. Then, as we did for the gene activity scores, we applied MAGIC to the TF z-score matrix to impute missing TF z-scores and smooth the TF z-scores across cells. The R implementation of MAGIC (Rmagic R package v2.0.3) was used and the algorithm was run with the function ‘magic’ using default parameters, except for the smoothing parameter, which was set to t=2. Finally, we downloaded TF families from the TFClass database (https://tffclass.bioinf.med.uni-goettingen.de)^33^ and computed TF family z-scores as the sum of z-scores of each TF in a given family divided by the square root of the number of TFs in the family.

#### Definition of cell clones with and without subclonal copy number changes of interest

Three high-frequency subclonal copy number changes of interest were identified from the copy number data: a gain in chromosome 5 and two losses in chromosomes 1 and X, respectively. To estimate their respective breakpoint positions, the copy number data was filtered as described in **Additional filtering of the copy number data**, except that the mappability constraint was relaxed to allow bins with mappability score of 0.9 and above. This ensured that conservative estimates of the breakpoints were obtained, by mostly removing genomic bins near centromeres and telomeres. For each copy number change, the breakpoints were estimated by the most frequent breakpoints observed across cells with the copy number change. These estimated breakpoints were then used to define two clusters of cells, respectively with, and without, the copy number change. The clustering procedure is illustrated using the loss in chromosome 1.

Cells that were diploid in the loss region defined by the estimated breakpoints, as well as adjacent regions of the same length, formed cluster C_D_. Cells that had copy number 1 in the loss region defined by the estimated breakpoints, and were diploid in the adjacent regions of the same length, formed cluster C_L_. In particular, cells with breakpoints that differed from the estimated breakpoints, and cells that did not have copy number equal to 2 or 1 in the entirety of the regions specified, were excluded. This procedure was applied similarly for the gain in chromosome 5 and the loss in chromosome X.

The three genomic regions of interest (in hg38 genome assembly) were then (1) 1-141,620,433 Mb on chromosome 5, with breakpoints between 45,499,898 Mb and 50,204,166 Mb and at 93,664,294 Mb; (2) 147,528,175-188,030,870 Mb on chromosome 1, with breakpoints at 161,030,210 Mb and 174,530,862 Mb, and (3) 1-114,256,790 Mb on chromosome X, with a breakpoint at 56,973,567 Mb. Of note is the uncertainty about the left-breakpoint of the gain on chromosome 5, which was situated to the left of the centromeric region in about half of the cells and to the right of the centromeric region in the other half of the cells. In this case, the breakpoint location was estimated as halfway between the two most frequent breakpoints and cells with either of these most frequent breakpoints were retained in the analysis.

#### Calculating ATAC signal in peaks for each copy number cluster

The ArchR project comprising all cells from our SW620 long-term chemotherapy exposure experiment was subset using the subsetArchRProject function to contain only the n=513 cells which passed the copy number QC (see ‘Additional filtering of the copy number data’). Dimensionality reduction was performed using the addIterativeLSI function, with iterations = 3, resolution = 1, and default parameters otherwise. Cells were then clustered using the addClusters function, with resolution = 0.8 and default parameters otherwise. These clusters were used to create pseudo-bulk replicates (addGroupCoverages function) and to call peaks using MACS2 (addReproduciblePeakSet function). The peak set was finally obtained using the getPeakSet function.

The copy number clusters defined in ‘Definition of cells clusters with and without subclonal copy number changes of interest’ were added into this ArchR project. Pseudo-bulk ATAC signal, normalised by reads in TSS, was obtained for each copy number cluster using the getGroupBW function. Combining the peak set with the ATAC signal for each copy number cluster, and restricting to the genomic regions of interest, the ATAC signal in peaks, and outside of peaks, was determined for each cluster. The 95^th^-percentile of the ATAC signal per bp outside of peaks was used to estimate cluster-specific background signal per bp. For each cluster, peaks with ATAC signal per bp smaller than or equal to the background ATAC signal per bp were excluded. This yielded a final peak set, with ATAC signal per peak, for each copy number cluster.

#### Comparing ATAC signal in peaks between copy number clusters

For each of the three copy number changes considered, the ATAC signal in the peaks shared by the two clusters of cells, respectively, with and without the copy number change was compared using the log2 fold change. For each shared peak, this was calculated as log2(signal in cluster with the change/signal in cluster without the change).

#### Testing for significant alterations in chromatin accessibility near breakpoints

For the three pairs of clones with and without CNAs, significant differences in mean log2 fold change (L2FC) in ATAC signal between the clones with and without the CNA, in regions flanking breakpoints, were tested for.

Excluding the first breakpoint on chromosome 5 due to its uncertainty about the centromere, 4 breakpoints were considered: 2 on chromosome 1, 1 on chromosome 5 and 1 on chromosome X. For each breakpoint, the left and right flanking regions were distinguished, and 1,000 random loci were sampled from the genomic segments (of constant copy number) to the left and right of the breakpoint, respectively. Then, the mean L2FC in ATAC signal within 100kb, 200kb, 500kb, 750kb, 1Mb and 1.5Mb to the left (or right) of the breakpoint and the corresponding random loci were calculated. Finally, the actual values of the mean L2FC in ATAC signal flanking the breakpoints were compared to the distributions of mean L2FCs in ATAC signal obtained using the random loci (i.e., the ‘null’ distributions). To determine whether the actual value was significantly lower (or higher) than the corresponding random distribution of values, a p-value was calculated as the percentile, P (or 100 – P), of the random distribution the actual value is equal to. The p-values were adjusted for multiple testing using the ‘p.adjust’ function from the stats R package v4.3.2, with the method set to “BH” for Benjamini-Hochberg correction^34^.

#### Processing the single-cell genotype calls from the WGS and ATAC data

The single-cell genotype calls from the WGS and ATAC data, stored in VCF files, were used to create SNP-by-cell matrices for both data modalities. These two matrices contain entries equal to 0, 1 and -1, which correspond to a SNP being, respectively, absent, present or unsampled in a cell.

In more detail, for each detected SNP position along the genome, cells were typically called to be in one of three states: ‘./.’, indicating a lack of reads at that position, ‘0/0’, indicating the cell has the reference allele at that position, and ‘1/1’ indicating the cell has the alternative allele at that position. We encoded these three states as -1, 0 and 1, respectively, in the SNP-by-cell matrices. A minority of cells were alternatively called as ‘0/1’ at a SNP position (< 2.4% of cells for 95% of SNP positions (WGS), < 1.25% of cells for all SNP positions (ATAC)), indicating that both alleles were sampled by chance. In these cases, we randomly sampled the state of the cell to be 0 or 1 (Bernoulli distribution, p = 0.5) in the SNP-by-cell matrices. Finally, for a minority of SNP positions along the genome (0.37% (WGS), 0.14% (ATAC)), we detected more than one alternative allele (e.g., some cells were called as ‘2/2’, ‘2/0’, ‘1/2’, in the case of 2 alternative alleles). In such cases, we defined distinct SNPs for each alternative allele. Given one alternative allele and the corresponding SNP, we encoded cells as 0 or 1 if there was evidence of the reference or this alternative allele, respectively, and cells with only evidence of another alternative allele were encoded as -1.

#### Defining high confidence heterozygous SNPs from the single-cell genotype calls from the WGS data

To identify heterozygous SNPs in the genomic regions of interest on chromosome 1 and 5, we first defined clusters of cells with a largely “unaltered” karyotype in these regions. For the loss in chromosome 1, we selected cells that were diploid for at least 90% of the 147-188 Mb region, and, for the gain in chromosome 5, we selected cells that had copy number 3 for at least 90% of the 1-47.5 Mb region and copy number 2 for at least 90% of the 47.5-141 Mb region. By treating these two cell clusters as pseudo-bulks, we then estimated the variant allele frequencies (VAFs) of each SNP in our WGS SNP-by-cell matrix as

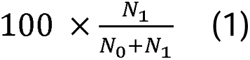

where N_0_ and N_l_ are the number of cells with the reference and alternative alleles, respectively, at the SNP position.

Heterozygous SNPs were then defined as the SNPs on chromosome 1 with VAFs between 40% and 60%, the SNPs on chromosome 5 within 1-47.5 Mb with VAFs between 23% and 43% or between 56% and 76%, and the SNPs on chromosome 5 within 47.5-141 Mb with VAFs between 40% and 60%. The final set of high confidence heterozygous SNPs was obtained by filtering for N_l_ > 10, i.e., at least 10 reads supporting the presence of each SNP.

We excluded the chromosome X region in this analysis as this chromosome was wholly duplicated prior to the loss at 1-57 Mb, implying the absence of heterozygous SNPs.

#### Computing heterozygous SNP VAFs in CNA clones from the WGS and ATAC data

We treated the previously defined CNA clones (C_L_ and C_D_ for the loss in chromosome 1 and C_G_ and C_D_ for the gain chromosome 5) as pseudo-bulks. Then, for each clone, we separately estimated SNP VAFs from the WGS and ATAC SNP-by-cell matrices using equation (1). For each clone, this yielded two VAF read-outs per SNP, one computed from the WGS data and one computed from the ATAC data.

#### Linear regression of the heterozygous SNP VAFs computed from the WGS data

Given the SNP VAFs computed from the WGS SNP-by-cell matrices for each CNA clone, we used the ‘lm’ function from the stats R package v4.3.2 to implement linear regression models predicting the absolute distance of SNP VAFs from 50% based on copy number, proximity to breakpoints and presence in open or closed chromatin region. Predictor variables were the copy number at the SNP position, multiplied by 10 (for visualisation purposes), and binary variables indicating whether a SNP was located within 1 Mb to the left or right of each breakpoint and whether a SNP was in a region of chromatin found to be open in the clone with the CNA or without the CNA.

For the loss in chromosome 1, this yielded 7 predictor variables, including 2 for SNPs in open chromatin regions in clone CL and clone CD, and 4 for SNPs located to the left and right of the two breakpoints. Due to the uncertainty in the breakpoint location around the centromere for the gain in chromosome 5, we only considered the 2^nd^ breakpoint, yielding 5 predictor variables in this case.

---

**Extended Data Figure 1.**
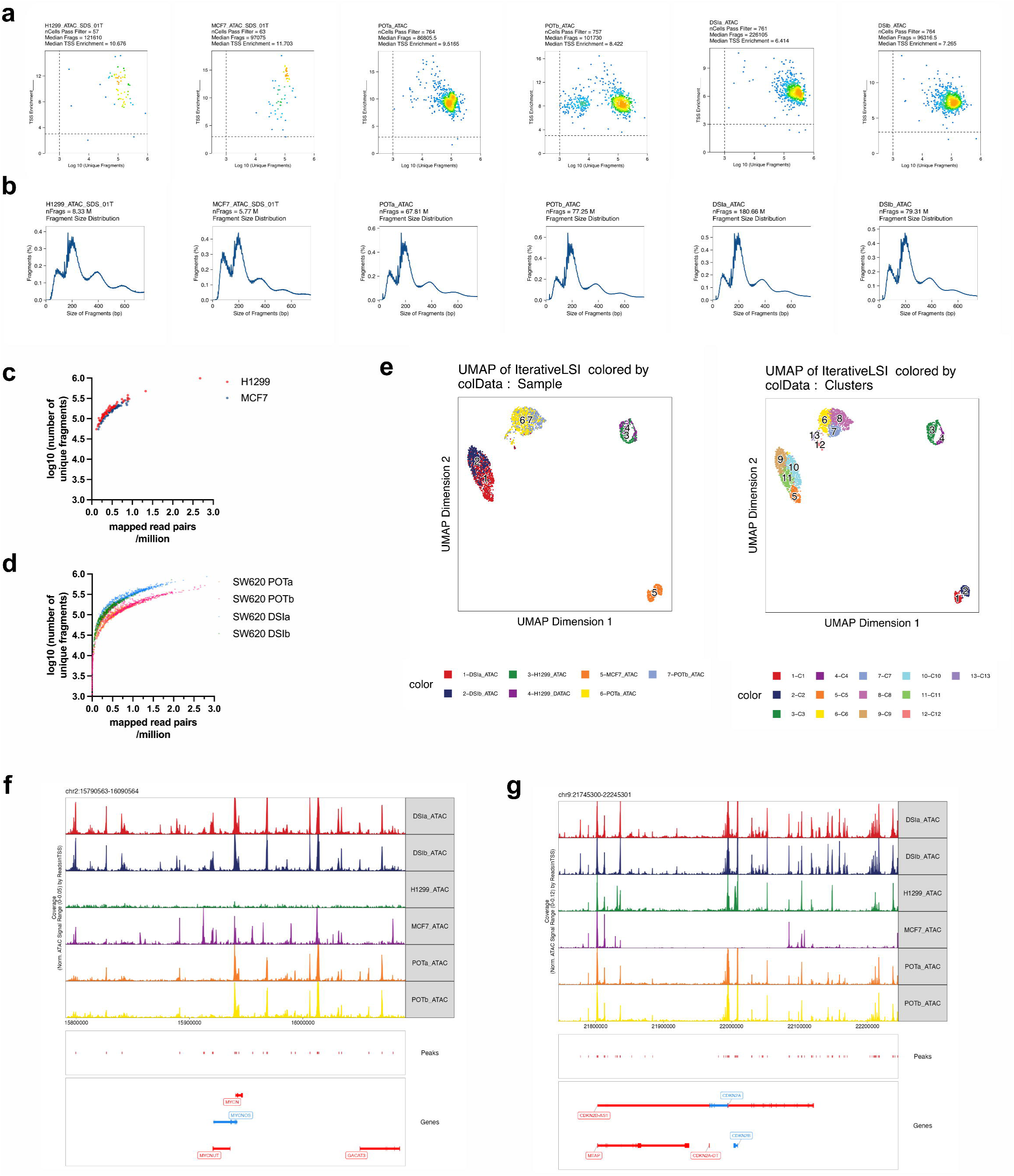
Technical performance of dATAC assay

**Extended Data Figure 2.**
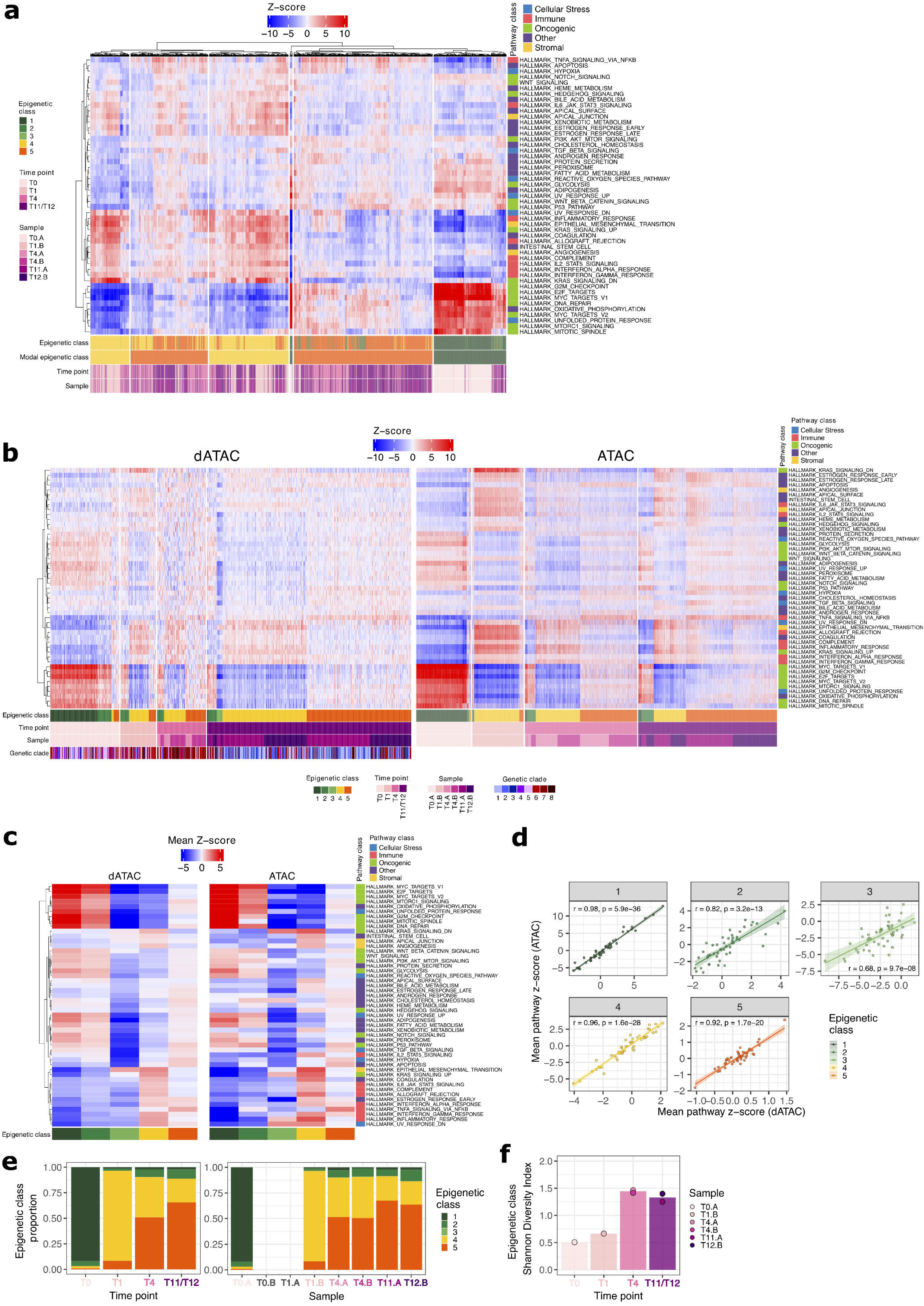
Validating the epigenetic classes defined from the dATAC data using the ATAC-only data. (a) Clustering of chromatin accessibility profiling for hallmark pathway loci from the ATAC-only data and classification of individual cells’ epigenetic class using the dATAC-derived classes divides ATAC-only cells into epigenetic clusters largely consistent with the 5 epigenetic classes defined from the dATAC data (Fig. 4d). (b) Comparison of hallmark pathway z-scores between dATAC and ATAC-only datasets, where individual cells were ordered by time and epigenetic class. Accessibility scores across time and epigenetic class were highly consistent between the two modalities. (c) Comparison of the mean hallmark pathway z-scores in each epigenetic class between dATAC and ATAC-only datasets further showed the concordance between the epigenetic profiles observed in both datasets. (d) The mean hallmark pathway z-scores in each epigenetic class, computed from the dATAC and ATAC-only datasets, were significantly correlated in all five classes. The lowest correlation was observed for class 3, but this class was significantly smaller than the others (9 cells (dATAC), 5 cells (ATAC)). (e) Epigenetic class sizes in the ATAC-only data evolved through 5-FU exposure (left panel) as in the dATAC data (Fig. 4e, left panel), although the ratio of class 5 to class 4 was larger than in the dATAC data at T11/T12. Very similar changes in epigenetic class size were also observed in the replicate evolution experiment (right panel). (f) Shannon diversity of epigenetic class by timepoint in the ATAC-only data showed increasing epigenetic class diversity after long-term 5-FU exposure.

**Extended Data Figure 3.**
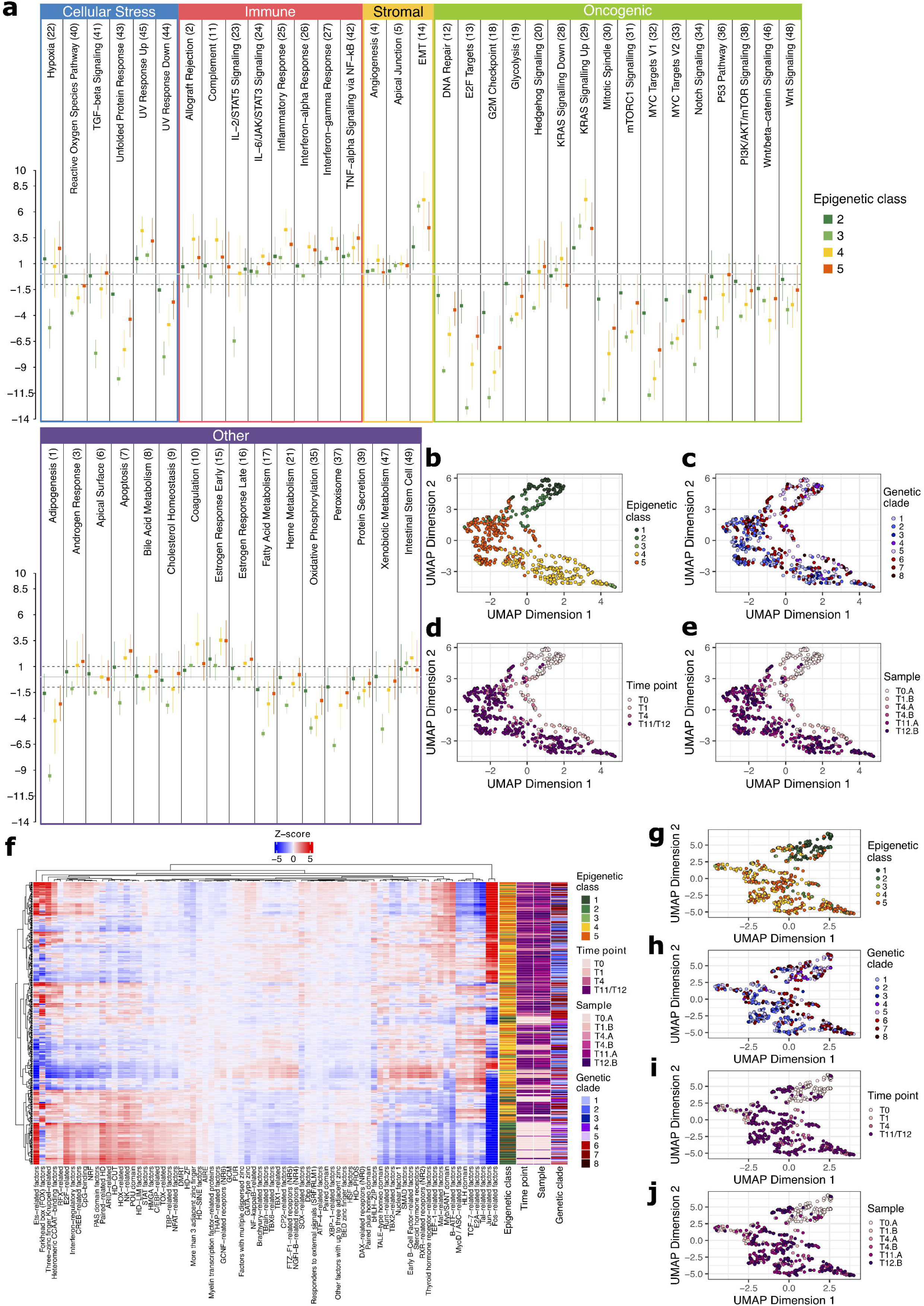
(a) Forest plot showing the mean and 95% confidence interval of hallmark pathway z-scores in epigenetic classes 2-5, normalised using the mean and standard deviation of hallmark pathway z-scores in epigenetic class 1 (all untreated cells bar one). The most significant differences in accessibility compared to class 1 were typically observed in classes 3 and 4, while class 2 was typically more like class 1. Class 5 appeared to be an intermediate state between classes 3,4 and class 2. (b-e) Hallmark pathway z-score UMAPs coloured by epigenetic class, genetic clade, time, and sample, show how treatment simultaneously causes a gradual (to class 2) and large (to classes 3 and 4) shift in epigenome, followed by stabilisation in the intermediate states (classes 4 and 5). (f) Heatmap of transcription factor (TF) family z-scores across individual cancer cells shows distinct TF activity profiles in untreated and treated cells. Most (> 80%) cells in drug-tolerance-associated epigenetic classes 4 and 5 had significantly higher accessibility of FOS and JUN-related TFs compared to untreated cells. (g-j) TF family z-score UMAPs coloured by epigenetic class, time, sample, and genetic clade, distinguish between untreated cells in epigenetic classed 1 and 2 and most treated cells in epigenetic classes 3-5.

**Extended Data Figure 4.**
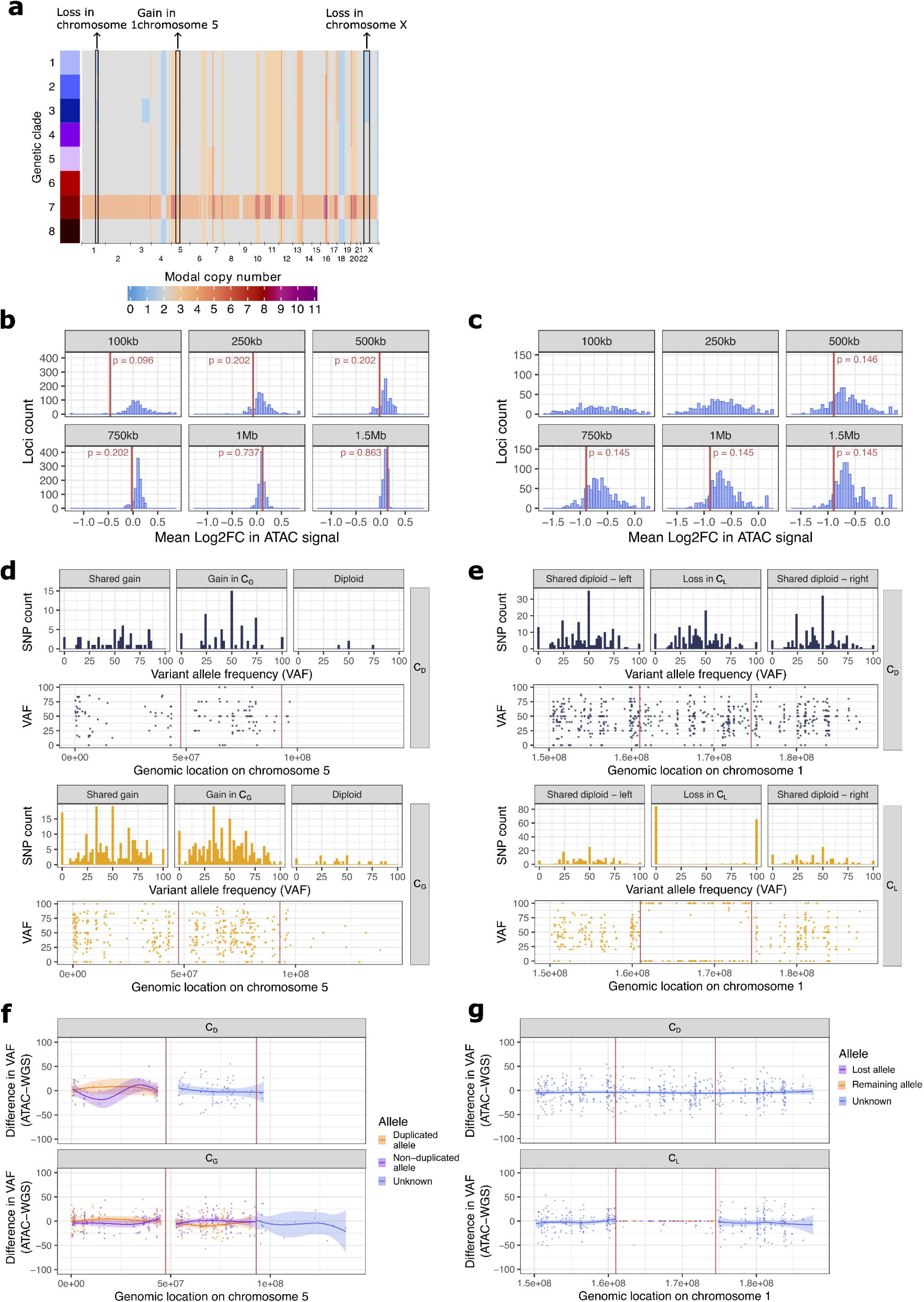
(a) Modal copy number in each genetic clade (from Fig 4a-b) highlighting three high-frequency subclonal copy number alterations: a gain in chromosome 5 and losses in chromosome 1 and X, respectively. (b) For the loss in chromosome 1, comparison of the ATAC signal in peaks between clones C_L_ and C_D_ within 100 kb – 1.5 Mb to the left of randomly selected loci located in the left common diploid region (blue histogram) and within 100 kb – 1.5 Mb to the left of the first breakpoint (red line). The p-values displayed are FDR-adjusted. (b) For the loss in chromosome X, comparison of the ATAC signal in peaks between clones C_L_ and C_D_ within 100 kb – 1.5 Mb to the left of randomly selected loci located in the loss region (blue histogram) and within 100 kb – 1.5 Mb to the left of the breakpoint (red line). There were no peaks within 250 kb of the breakpoint in the loss region, hence the lack of true value displayed for 100 kb and 250 kb. The p-values displayed are FDR-adjusted. (d-e) VAFs of heterozygous SNPs in the regions of interest on chromosomes 5 and 1, computed from the ATAC component of the dATAC data. The VAF distributions are consistent with those in Fig. 5e,f, though noisier due to lower read depth. (f) For clones C_G_ and C_D_, differences in heterozygous SNP VAFs computed using the ATAC and WGS components of the dATAC data were plotted by location on the chromosome 5 region of interest. Differences were equally distributed about 0 along the entire segment, supporting the assumption of equal allele accessibility. (g) For clones C_L_ and C_D_, differences in heterozygous SNP VAFs computed using the ATAC and WGS components of the dATAC data were plotted by location on the chromosome 1 region of interest. As in (f), the plots support the assumption of equal allele accessibility.

